# Multifaceted control of E-cadherin dynamics by the Adaptor Protein Complex 1 during epithelial morphogenesis

**DOI:** 10.1101/2021.11.08.467679

**Authors:** Miguel Ramírez Moreno, Katy Boswell, Helen L. Casbolt, Natalia A. Bulgakova

## Abstract

Intracellular trafficking regulates the distribution of transmembrane proteins including the key determinants of epithelial polarity and adhesion. The Adaptor Protein 1 (AP-1) complex is the key regulator of vesicle sorting, which binds many specific cargos. We examined roles of the AP-1 complex in epithelial morphogenesis, using the *Drosophila* wing as a paradigm. We found that AP-1 knockdown leads to ectopic tissue folding, which is consistent with the observed defects in integrin targeting to the basal cell-extracellular matrix adhesion sites. This occurs concurrently with an integrin-independent induction of cell death, which counteracts elevated proliferation and prevents hyperplasia. We discovered a distinct pool of AP-1, which localizes at the subapical Adherens Junctions. Upon AP-1 knockdown, E-cadherin is hyperinternalized from these junctions and becomes enriched at the Golgi and recycling endosomes. We then provide evidence that E-cadherin hyperinternalization acts upstream of cell death in a potential tumour-suppressive mechanism. Simultaneously, cells compensate for elevated internalization of E-cadherin by increasing its expression to maintain cell-cell adhesion.

**Author Summary:** The epithelium is one of the four types of tissues found in animals and is essential for the normal development and maintenance of multicellular organisms. In this tissue, the adhesion protein E-cadherin helps keep cells together and facilitates the coordination of their behaviours. E-cadherin is highly dynamic and undergoes constant turnover by intracellular trafficking machinery. Here, we describe the contributions of one of the core components of intracellular trafficking, the AP-1 complex to E-cadherin dynamics. Using epithelial cells from *Drosophila melanogaster*, we discover that the AP-1 complex limits E-cadherin internalization from the plasma membrane, which is consistent with the localization of a distinct pool of the AP-1 complex at the sites of E-cadherin cell-cell adhesion. We also show that increased E-cadherin internalization triggers programmed cell death, preventing the tissue from hyperplastic overgrowth in a potential tumour-suppressive mechanism. At the same time, cells compensate for the reduction in the membrane presentation of E-cadherin by increasing its expression, therefore protecting tissue integrity.

## Introduction

Epithelial morphogenesis gives origin to complex structures from simple epithelial sheets [1]. It is driven by changes in the properties of the cells, often influenced by adhesion molecules [1,2]. These transmembrane molecules constitute the cornerstone of interactions between individual cells and with their environment, enabling morphogenetic processes [3,4]. E-cadherin (E-cad, encoded by the *shotgun* gene in *Drosophila*) is one of the cell-cell adhesion molecules that connect neighbouring cells at the subapical Adherens Junctions (AJs) in epithelial cells [5,6]. E-cad contributes to numerous developmental processes and diseases such as cancer [3,7,8]. Basally, integrins anchor cells to the extracellular matrix, controlling tissue folding but also linking the cell architecture and intracellular signalling [9–12]. The regulation of the cell surface presentation of these adhesion proteins and their impact on tissue morphogenesis remain open questions in the fields of cell and developmental biology [13].

Intracellular trafficking represents one of the main ways to regulate protein presentation at the cell surface [14,15]. Cargo-containing vesicles are sent from and to numerous organelles, with the Trans Golgi Network (TGN) and recycling endosomes (REs) serving as the main sorting centres in the cell [15–17]. Most of what is known about trafficking pathways and their key regulators come from studies in single cells, which yield a great molecular data output but do not inform about their functions in tissue contexts during development [7,18–21].

One of the key regulators of intracellular trafficking is the Adaptor Protein Complex 1 (AP-1), shuttling vesicles within the TGN/REs continuum [15,16,22,23]. It is a heterotetrameric complex consisting of two large subunits (γ and β), one medium subunit (μ) and one small subunit (σ)[24]. Two mammalian isoforms exist, depending on the participant μ subunit: the μ1-containing ubiquitously expressed AP-1A and the μ2-containing epithelia-specific AP-1B [25–28]. Although the mammalian μ subunits are 79% identical, AP-1A sorts basolateral cargos at the TGN, where it is recruited by the small GTPase Arf1 [27,29–31] and AP-1B sorts proteins from REs to the basolateral membrane and is regulated by the small GTPase Arf6 (Shteyn et al., 2011). AP-1B is also found at integrin-mediated focal adhesions, and its loss correlates with the migratory behaviour of metastatic cells (Kell *et al*., 2020). Two AP-1s are also found in *Caenorhabditis elegans*, where they restrict the basolateral localization of E-cad [32,33]. In the fruit fly, *Drosophila melanogaster*, a single AP-1 complex localizes to both TGN and REs and is required for the maintenance of E-cad at the ring canals during oogenesis [34–36]. In epithelia, the fly AP-1 has only been linked to the distribution of Notch pathway components in specialized cells – the retina and sensory organs [34,35,37]. There, it is one of the pathways, which modulates the trafficking of Notch towards E-cad junctions [35,38]. Curiously, more Notch accumulates at these junctions upon loss of AP-1, which was attributed to elevated apical targeting at the expense of basolateral without apparent defects in cell polarity and E-cad adhesion.

In this work, we demonstrate that the *Drosophila* AP-1 complex is necessary for epithelial morphogenesis using the developing wing as a paradigm. There, the AP-1 complex influences cell size and number, and tissue folding and controls the transport of E-cad and integrins. We identified a distinct subapical pool of the AP-1 complex and determined that the AP-1 complex modulates the endocytosis of E-cad. The excessive internalization of E-cad following AP-1 knockdown triggers cell death, which counteracts elevated proliferation and suppresses hyperplasia: overgrowth without disrupting the epithelial architecture [39]. The surface levels of E-cad remain however stable due to adjusted expression. Altogether, our results demonstrate the versatility of functions of the AP-1 complex in intracellular trafficking in vivo and link it to tissue development and pathology.

## Results

We employed the GAL4/UAS system [40] and the *engrailed* promoter (*en::*GAL4) to knockdown AP-1 using RNAi throughout development in the posterior compartments of the wing primordia (imaginal discs, Fig 1A-C). Downregulation of AP-1μ or AP-1γ (Fig S1A) – two unique subunits of the AP-1 complex [16,41] – led to defective morphology of adult wings, ranging from absent to severely disrupted posterior compartments (Fig 1A). A similar adult phenotype was caused by using *MS1096*::GAL4 driver (Fig S1B), which expresses GAL4 throughout the wing pouch (presumptive region of the adult wing, Fig 1C).

**Figure 1.**
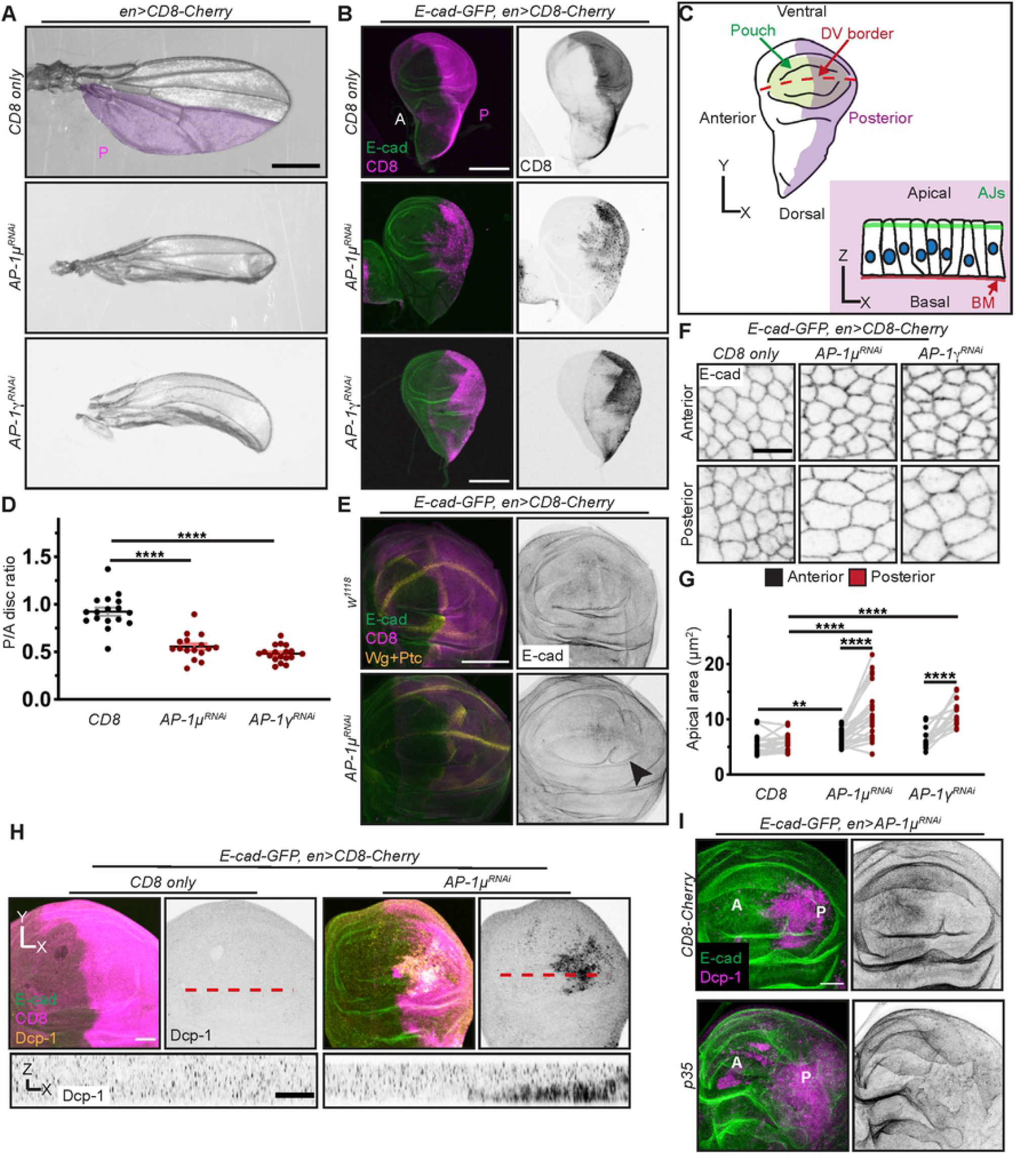
AP-1 regulates development, cell morphology and survival in the wing disc. (A) Wings expressing CD8-Cherry alone or with AP-1 RNAis in the posterior compartment. Scale bar: 0.5 mm. (B) Wing discs expressing CD8-Cherry (magenta, left; grayscale, right) alone or with AP-1 RNAis, showing E-cad-GFP (green, left), and Anterior (A) and Posterior (P) compartments. Scale bar: 150 μm. (C) Top – cartoon view of a wing imaginal disc with posterior compartment (magenta), wing pouch (green) and dorsoventral border (DV, dashed red line). Bottom – sagittal view of the wing pouch epithelium with the AJs (green) and the basal membrane (BM, red). (D) Posterior:Anterior (P/A) area ratios of control and AP-1 RNAis expressing discs. Dots represent individual discs (n=17, 16 and 18). ****P<0.0001 (Brown-Forsythe and Welch ANOVA test). (E) Wing pouches expressing CD8-Cherry alone or with AP-1μ RNAi, co-stained for Wingless and Patched (Wg and Ptc, both in yellow). E-cad-GFP visualizes the tissue architecture (green, left; grayscale, right). Arrowhead indicates an ectopic fold. Scale bar: 100 μm. (F) Dorsal wing pouch cells expressing *shg*::E-cad-GFP (grayscale) in the indicated compartments (rows) and genotypes (columns). Scale bar: 5 μm. (G) Cell apical area of wing pouch cells. Each dot represents the average of an individual disc (n=15-23 discs/genotype). **P<0.01 and ****P<0.0001 (Wilcoxon and paired t-test, between compartments; and Kluskal-Walls or One-Way ANOVA tests, between genotypes). (H) Wing pouches expressing CD8-Cherry alone or with AP-1μ RNAi with E-cad-GFP (green, left), CD8-Cherry (magenta, left) and cleaved effector caspase (Dcp-1, yellow, left; grayscale, right and bottom sagittal projection of the dashed line). Scale bars: 50 μm (top) and 20 μm (bottom). (I) Wing pouch regions of discs expressing AP-1μ RNAi with CD8-Cherry (top, not shown) or p35 (bottom) in the posterior compartment visualized with E-cad-GFP (green, left; inverted grayscale, right) and cleaved caspase-3 (DCP-1, magenta, left). Anterior (A) and Posterior (P) compartments are indicated. Scale bar: 50 μm.

We observed complex morphological changes upon AP-1μ or AP-1γ knockdown already in third instar wing imaginal discs (Fig 1B-E). Posterior compartments were reduced in size, as reflected by their Posterior/Anterior (P/A) compartment ratio (Fig 1B, D). AP-1 knockdown resulted in ectopic folding of the wing pouch region, although compartmental borders remained well defined in the disc (Fig 1E). While this folding could contribute to the reduced compartment size, we also examined the apical cell area and found that it was increased, as detected with GFP-tagged E-cad expressed from the endogenous locus (the knock-in of GFP into *shg* gene, *shg*::E-cad-GFP, Fig 1F-G [42]). Combining the apical cell area increase with the reduced size of compartments indicated a decrease in the total number of cells in the monolayer epithelium of the wing pouch. This could be a consequence of a reduced proliferation rate, increase in cell death, or extrusion of live cells. However, the proliferation rate was increased when normalized to cell number, although the rate on the tissue level was unaffected (Fig S1C-E). Concurrently, we detected considerable cell death visualised by the cleaved effector caspase Dcp-1 (Fig 1H, Fig S1F), which was accompanied by fragmented DAPI staining, indicative of nuclear debris (Fig S1G). The basal localization of Dcp-1 and fragmented DAPI (Fig 1H and Fig S1F, G) suggests that these are dying cells being eliminated and extruded [43,44]. Consistent with these changes in cell proliferation and death, co-expression of AP-1μ RNAi with the apoptosis-blocking baculoviral p35 protein, which blocks effector caspases but not their processing [45–47], caused hyperplastic growth (Fig 1I). Therefore, the three main effects of AP-1 knockdown in wing imaginal discs are increased cell death, enlarged apical cell area, and ectopic folds.

As ectopic folds can be due to altered basal adhesion [12], we examined integrin localization upon AP-1 knockdown. The αPS1 (Mew) integrin – specific for the dorsal compartment [48] – was nearly absent from the basal surface of the tissue (Fig 2A). We found similar changes in the distribution of αPS1 binding partner, βPS (Mys) and Laminin B2 (Fig S2A), further supporting the loss of basal adhesions. Mimicking this loss by overexpressing diβ, a dominant negative version of Mys which displaces endogenous protein (Fig 2B) [10,49] led to ectopic folds similar to AP-1 knockdown (Fig 2C) [10]. This correlated with the absence of αPS1 at the posterior compartment (Fig 2D), akin to the effects of AP-1 knockdown (Fig 2A). However, diβ expression did not increase the apical cell area (Fig 2E, F) and caused only small pockets of dying cells (Fig 2G) in contrast to AP-1 knockdown. These results are consistent with AP-1 downregulation leading to ectopic folding due to disruption of the basal localization of integrins. However, the effects of AP-1 downregulation on cell survival appear to be through an integrin-independent mechanism.

**Figure 2.**
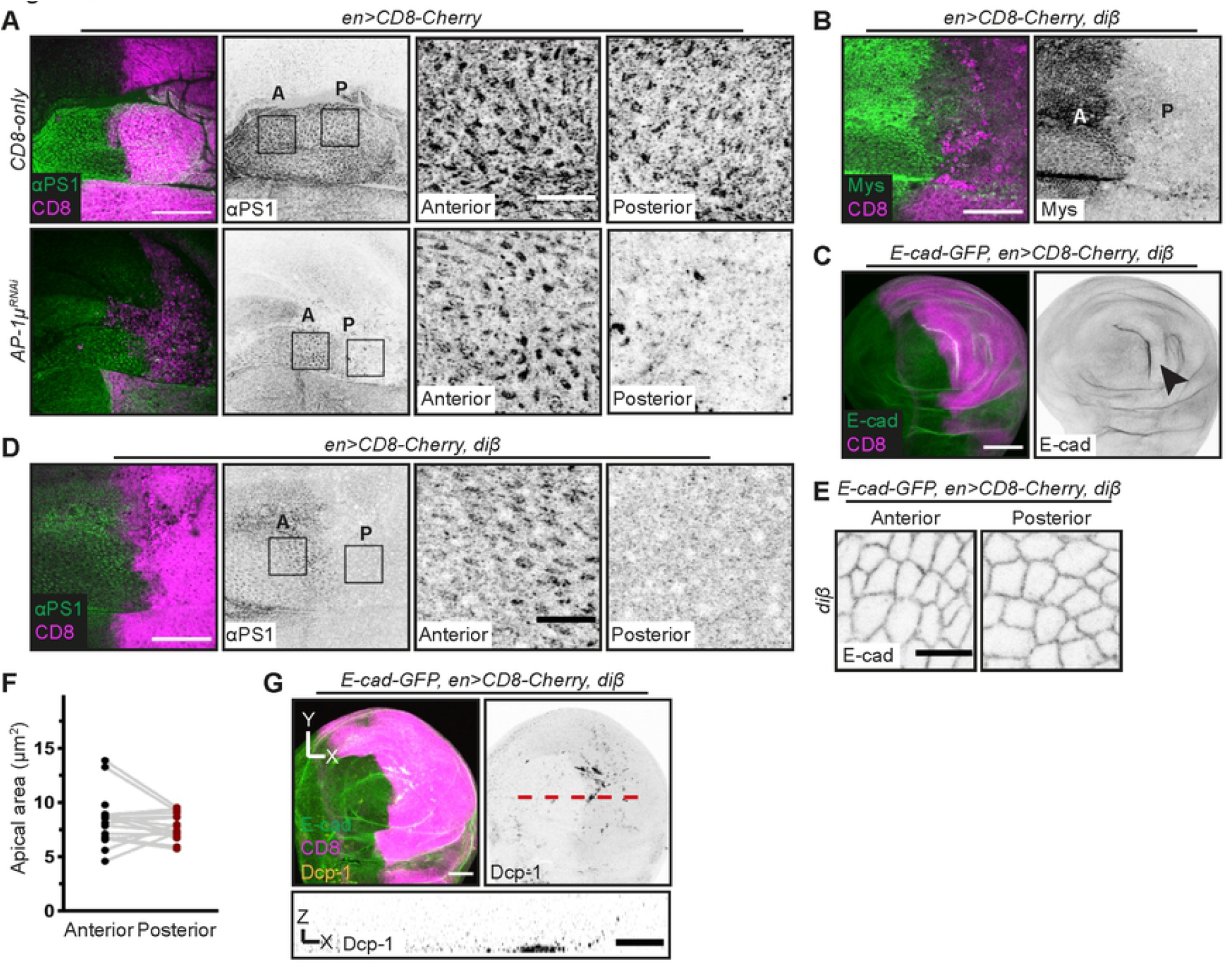
Disruption of basal polarity is not responsible for the AP-1 knockdown phenotype. (A) Basal region of wing discs expressing CD8-Cherry alone or with AP-1μ RNAi, co-stained with αPS1 (green, left; grayscale, right). Anterior (A) and Posterior (P) compartment areas (right panels) are highlighted by rectangles (left). Scale bars: 50 μm (left); 10 μm (right). (B) Basal region n of wing disc expressing CD8-Cherry (magenta, left) and diβ, stained for βPS integrin (Mys, green, left; inverted grayscale, right) and confirming the displacement of the endogenous βPS. Areas in the Anterior (A) and Posterior (P) compartments are indicated. Scale bar: 50 μm. (C) Wing pouch regions of discs expressing CD8-Cherry protein (magenta, left) with diβ in the posterior compartment. E-cad-GFP is shown to visualize tissue architecture (green, left; inverted grayscale, right). Arrowhead indicates an ectopic fold. Scale bar: 100 μm. (D) Basal region of wing discs expressing CD8-Cherry with diβ, stained with αPS1 integrin (green, left; grayscale, right). Areas in the Anterior (A) and Posterior (P) compartments (right panels) are highlighted by rectangles (left). Scale bars: 50 μm (left); 10 μm (right). (E) Dorsal wing pouch cells expressing *shg*::E-cad-GFP (grayscale) expressing CD8-Cherry with diβ in the posterior compartment. Scale bar: 5 μm. (F) Cell apical area of discs depicted in E. Each dot represents the average of an individual disc; compartments from same disc are connected by grey lines (n=22 discs). (G) Wing pouches expressing CD8-Cherry with diβ, visualised with E-cad-GFP (green, left), CD8-Cherry (magenta, left) and cleaved effector caspase (Dcp-1, yellow, left; grayscale, right and bottom sagittal projection of the dashed line). Scale bars: 50 μm (top) and 20 μm (bottom).

To gain insights into this mechanism, we explored the intracellular localization of the AP-1 complex using the Venus-tagged AP-1μ (AP-1μ-VFP) [35]. Consistent with other cell types [15,16,35,36], AP-1μ-VFP accumulated within puncta (spots) matching the TGN (Golgin-245-positive, [50,51], Fig 3A) and REs (Rab11-positive, Fig 3A, [52]). Unexpectedly, we also found a distinct subapical pool of AP-1μ colocalizing with E-cad at the wing discs AJs (Fig 3A-C), as well as in the embryonic epidermis and pupal eyes (Fig S3A-B). In contrast, we detected only a few puncta at the basal surface, with no αPS1 co-localization (Fig 3A and Fig S3C). AP-1γ knockdown reduced levels of AP-1μ-VFP in the cytoplasm and at AJs (Fig 3D-F), which we measured as either the mean intensity of native VFP fluorescence or its total content to account for the increased apical perimeter. These results suggested that the AP-1μ-VFP assembles into the AP-1 complex at AJs as well as in the cytoplasm. Indeed, we observed a co-localization at AJs between E-cad-GFP and an AP-1γ antibody (Fig S3D) [35].

**Figure 3.**
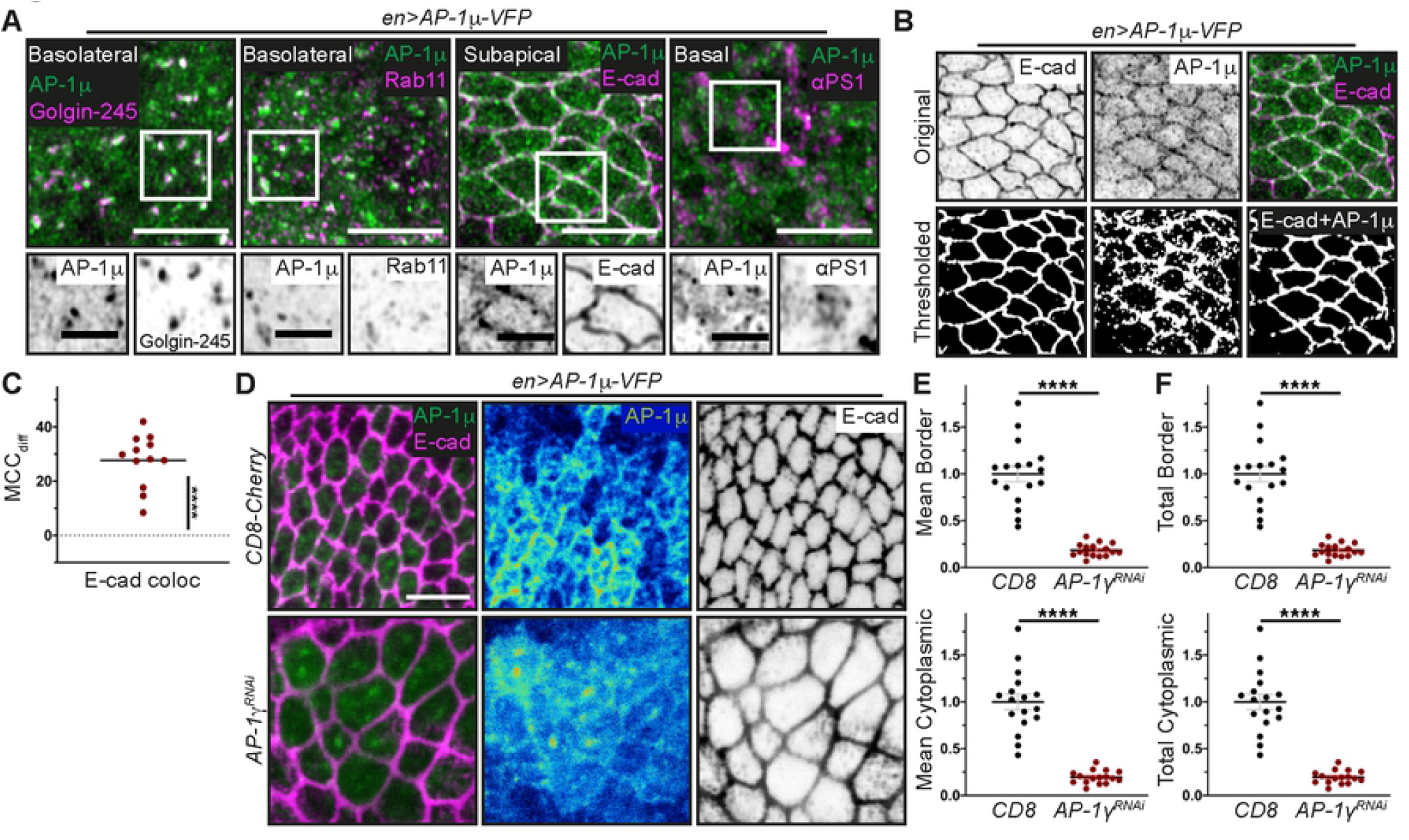
AP- 1 displays an apical pool colocalizing with E-cadherin. (A) Posterior compartments of wing discs expressing AP-1μ-VFP (green) and co-stained with markers (magenta) of the TGN (Golgin-245), RE (Rab11), the AJs (E-cad) and the basal membrane (integrin αPS1). Grayscale images correspond to AP-1μ-VFP and each marker within the white square on the image above. Scale bars: 5 μm (top) or 2 μm (detail). (B) Detail of the colocalization between E-cad antibody (inverted grayscale, top left; magenta, top right) and AP-1μ-VFP (inverted grayscale, top centre; green, top right), as shown originally in A. Bottom panels depict automatic thresholded images (left and centre) and the pixels that are positive for both thresholded channels (bottom right). (C) Manders’ Correlation Coefficient difference (bottom, MCC_diff_) for the E-cad and AP-1μ-VFP signals (n=12). ****P<0.0001 (one sample t-test in comparison to zero for random colocalization). (D) Posterior compartments of wing discs expressing AP-1μ-VFP (green, left; heatmap, right) together with CD8-Cherry (not shown, top) or AP-1γ RNAi (bottom) co-stained for E-cad (magenta, left; grayscale, right). Scale bar: 5 μm. (E, F) Mean (E) and total (F) AP-1μ-VFP levels at the cell borders (top) and in the cytoplasm (bottom). Each dot represents average for an individual disc (n=17, 16). ****P<0.0001 (two-tailed t-test).

To explore the roles of the AP-1 complex at AJs, we analysed the distribution and levels of E-cad. The mean levels of *shg*::E-cad-GFP at AJs were mildly reduced, while the total junctional protein content was unchanged or mildly elevated following AP-1 knockdown (Fig 4A-C). At the same time, we observed intracellular vesicles containing *shg*::E-cad-GFP, which are rarely seen in the control (Fig 4A-B). To examine whether AP-1 knockdown led to the accumulation of E-cad intracellularly, we measured the cytoplasmic intensity of *shg*::E-cad-GFP in the plane of AJs (“subapical cytoplasm” hereafter). Although it does not include all of the cytoplasm, both TGN and REs are found in this region (Fig S4) and the signal can be confidently attributed to individual cells. The mean levels of *shg::*E-cad-GFP in the subapical cytoplasm were also mildly reduced (Fig 4C). However, when normalized to the increase in the apical cell area, the total subapical intracellular content of *shg::*E-cad-GFP was elevated in agreement with the observed increase of E-cad containing puncta (Fig 4D). Therefore, AP-1 knockdown elevated the ratio of *shg::*E-cad-GFP content in the subapical cytoplasm relative to the plasma membrane (Fig 4E). Additionally, an increase in the cytoplasmic content combined with no change in the junctional *shg::*E-cad-GFP may indicate an increase in the total amount of protein per cell, which could be either due to elevated protein production or reduced degradation. We tested if there were changes in the protein production and found that AP-1 knockdown indeed increased *shg* mRNA levels (Fig 4F).

**Figure 4.**
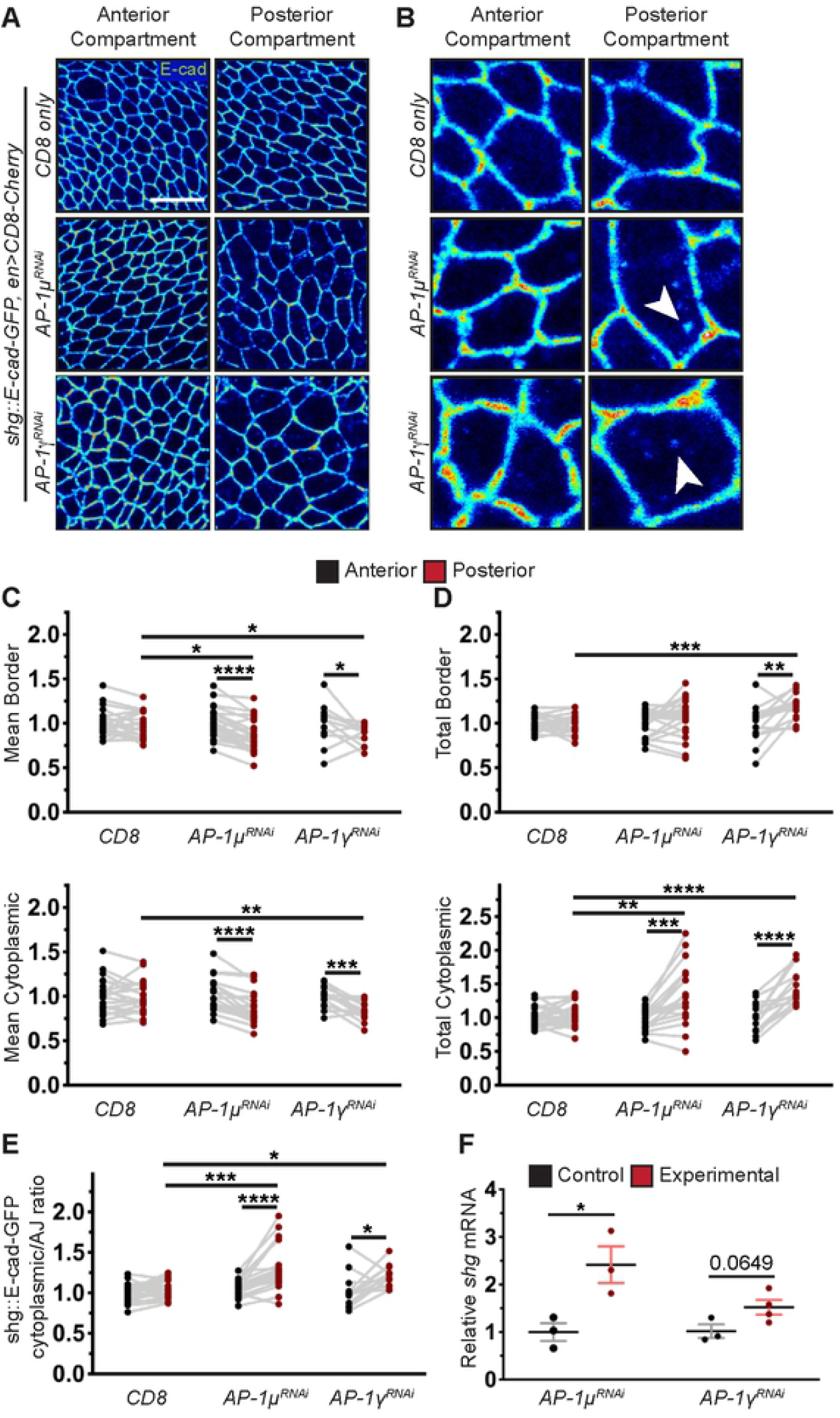
AP-1 controls the levels and expression of E-cadherin. (A) Dorsal wing disc pouch cells expressing *shg*::E-cad-GFP (heatmap) in the indicated compartments (columns) and genotypes (rows). Scale bar: 10 μm. (B) Detail of the panels in A. White arrowheads indicate intracellular E-cad-GFP accumulation. (C, D) Mean (C) and total (D) *shg*::E-cad-GFP protein levels at the cell borders (top) and in the cytoplasm (bottom). (E) Cytoplasm/AJs ratio of *shg*::E-cad-GFP total levels. (F) *shg* expression levels in control and with AP-1 subunits knockdown. On all graphs, dots represent individual discs (n= 22, 23, 15), apart from F, where each dot is an independent biological replicate (n= 3,3,3 and 4 per genotype); data from same discs are connected by grey lines. *P<0.05, **P<0.01, ***P<0.001 and ****P<0.0001. Wilcoxon and paired t-test (between compartments) and Kluskal-Walls or One-Way ANOVA tests (between genotypes) were used in C-E; Welch’s t-test in F.

To exclude the effects of AP-1 knockdown on *shg* gene expression, we used the GFP-tagged E-cad expressed from the ubiquitous *Ubi-p63E* promoter (*ubi*::E-cad-GFP, Fig 5A [53]). The levels of *ubi*::E-cad-GFP are similar to that of *shg::*E-cad-GFP (Fig 5E-F, Fig S5), suggesting physiological levels of E-cad in these cells in the control. We confirmed that the levels of *ubi*::E-cad-GFP protein were insensitive to *shg* mRNA expression by measuring them in the presence of the loss of function s*hg*^*IG27*^ allele [54]. In agreement with our recent findings of a feedback loop that stabilizes the levels of E-cad in wing discs [55], levels of *shg::*E-cad-GFP increased to compensate for the absence of the second gene copy (Fig 5E, F, Fig S5). However, levels of *ubi*::E-cad-GFP were not affected by the s*hg*^*IG27*^ allele (Fig 5E, F, Fig S5). At the same time, we observed that the downregulation of both μ and γ AP-1 subunits halved the *ubi*::E-cad-GFP levels at the AJs (Fig 5A-C) [56,57]). Concurrently, while the mean signal of *ubi*::E-cad-GFP in the subapical cytoplasm was also reduced, the total subapical intracellular content remained normal as a consequence of the increased apical cell area (Fig 5B-C). Therefore, for *ubi*::E-cad-GFP, AP-1 knockdown also elevated the ratio of its content in the subapical cytoplasm relative to the plasma membrane (Fig 5D).

**Figure 5.**
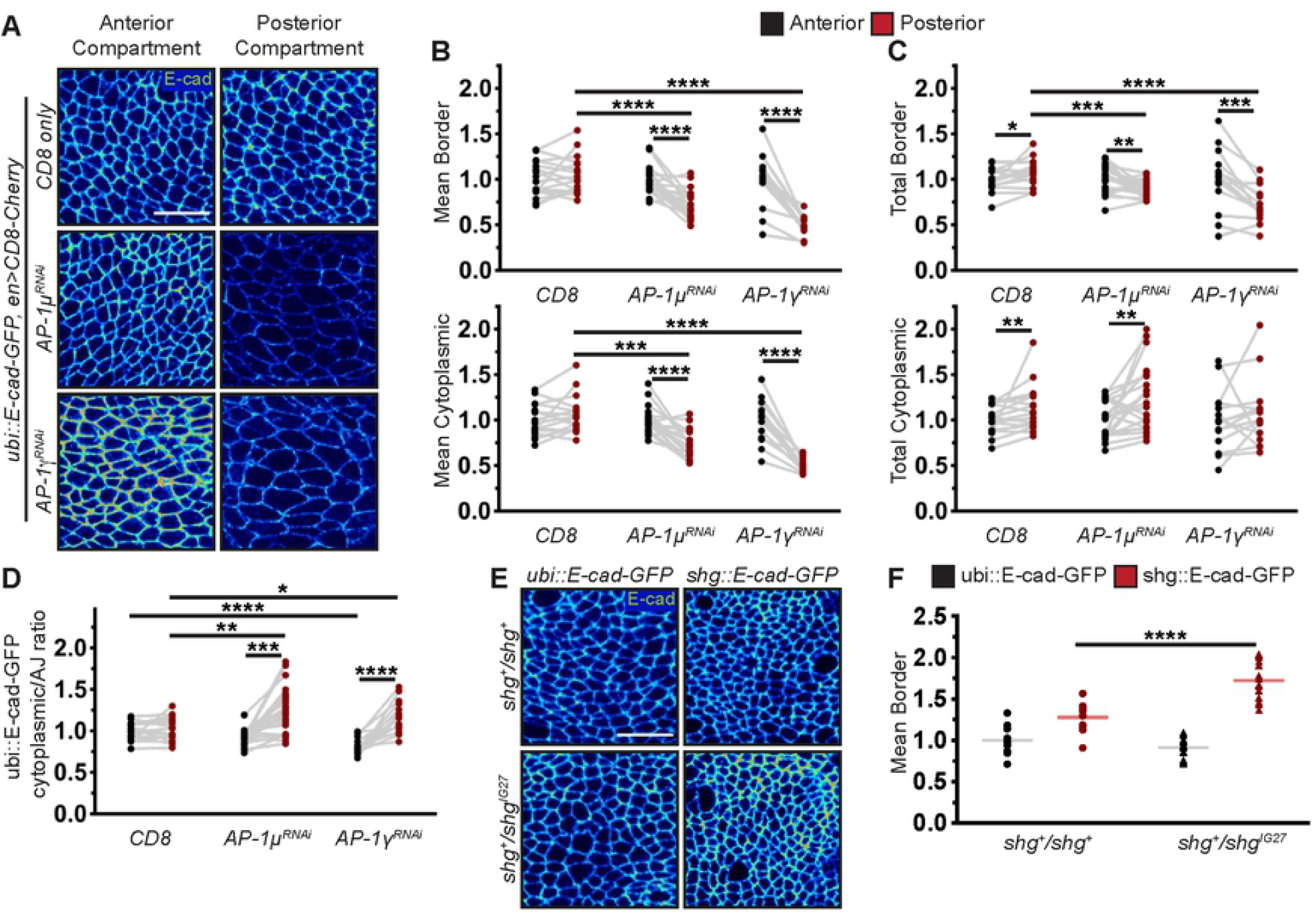
AP-1 controls the levels and expression of E-cadherin (cont.) (A) Dorsal wing pouch cells expressing *ubi*::E-cad-GFP (heatmap) in the indicated compartments (columns) and genotypes (rows). Scale bar: 10 μm. (B, C) Mean (B) and total (C) *ubi*::E-cad-GFP levels at the cell borders (top) and in the cytoplasm (bottom). (D) Cytoplasm/AJs ratio of the total levels for *ubi*::E-cad-GFP. (E) Dorsal wing pouch cells expressing E-cad-GFP (heatmap) expressed with the indicated constructs (columns) in the different genotypes (rows). Scale bar: 10 μm. (F) Mean E-cad-GFP levels at the cell borders as depicted in E. On all graphs, dots represent individual discs, (n= 17, 23, 15 in B, C and 14, 13, 14, 14 in F); data from same discs are connected by grey lines in B and C. *P<0.05, **P<0.01, ***P<0.001 and ****P<0.0001. Wilcoxon and paired t-test (between compartments) and Kluskal-Walls or One-Way ANOVA tests (between genotypes) were used in B, C, E, F; unpaired t-test in G.

The increase in the relative level of E-cad in the cytoplasm at the level of AJs and the intracellular puncta observed for *shg::*E-cad-GFP indicated E-cad accumulation in one or several intracellular compartments. Due to the localization of the AP-1 complex at the TGN and REs, we examined colocalization of both E-cad-GFP proteins with these compartments in the subapical cytoplasm and found that AP-1μ knockdown increased the colocalization in all cases (Fig 6A-F). We also found a lack of co-localization of E-cad-GFP proteins with the Golgi bodies in the controls, which is indicative of a very low residency time of E-cad-GFP there in the control. The colocalization was, however, noticeable, and significantly above zero following AP-1 knockdown (Fig 6C, F).

**Figure 6.**
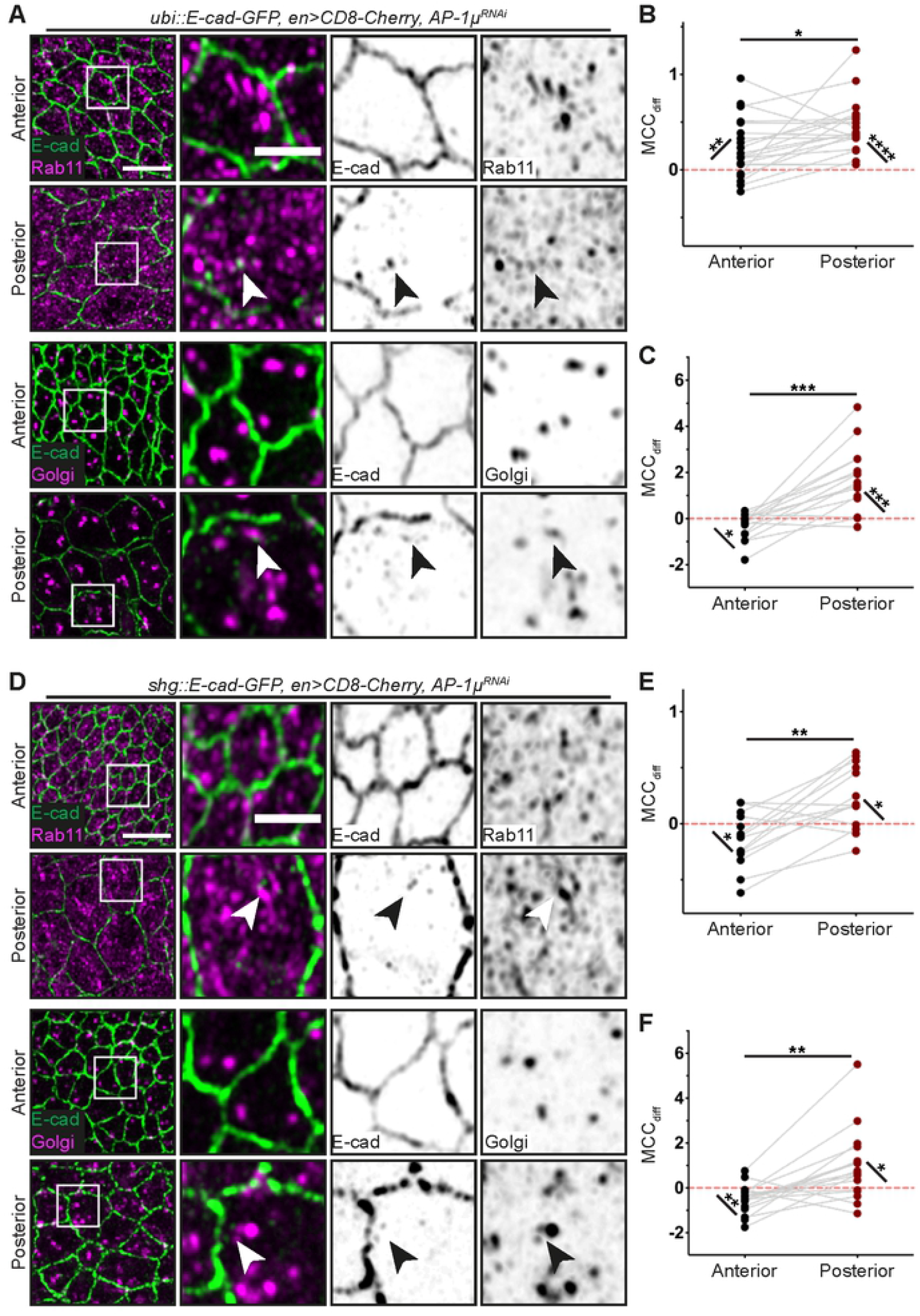
AP-1 knockdown increases the intracellular pool of E-cadherin. (A) Dorsal wing pouch regions of discs co-expressing *ubi::E-cad-GFP* (green) with CD8-Cherry (not shown) and AP-1μ RNAi in the posterior compartment. Discs are stained for Rab11 (top two rows) and Golgi mab (bottom two rows), both depicted in magenta. Right panels show magnified regions within the white squares in left panels. Arrowheads indicate co-localization of E-cad and each marker. Scale bars: 5 μm (left) and 2 μm (detail). (B) Manders’ Correlation Coefficient difference (MCC_diff_, see Methods) for the co-localization between Rab11 and *ubi::E-cad-GFP*. Each dot represents average for individual paired compartments (n=20 discs). *P<0.05 (Wilcoxon test); **P<0.01 and ***P<0.001 (one sample t-test). (C) Manders’ Correlation Coefficient difference (MCC_diff_, see Methods) for the co-localization between Golgi and *ubi::E-cad-GFP*. Each dot represents average for individual paired compartments (n=15 discs). ***P<0.05 (Wilcoxon test and one sample t-test); *P<0.01 and ***P<0.001 (one sample t-test). (D) Dorsal wing pouch regions of discs co-expressing *shg::E-cad-GFP* (green) with CD8-Cherry (not shown) and AP-1μ RNAi in the posterior compartment. Discs are stained for Rab11 (top two rows) and Golgi mab (bottom two rows), both depicted in magenta. Right panels show magnified regions within the white squares in left panels. Arrowheads indicate co-localization of E-cad and each marker. Scale bars: 5 μm (left) and 2 μm (detail). (E) Manders’ Correlation Coefficient difference (MCC_diff_, see Methods) for the co-localization between Rab11 and *shg::E-cad-GFP*. Each dot represents average for individual paired compartments (n=14 discs). *P<0.05 (one sample t-test) and **P<0.01 (Wilcoxon test). (F) Manders’ Correlation Coefficient difference (MCC_diff_, see Methods) for the co-localization between Golgi and *shg::E-cad-GFP*. Each dot represents average for individual paired compartments (n=17 discs). *P<0.05 (one sample t-test) and **P<0.01 (Wilcoxon test and one sample t-test).

A decrease in the plasma membrane levels and increase of the relative intracellular accumulation of *ubi*::E-cad-GFP could be either due to elevated protein internalization from the surface, its retention within intracellular compartments or a combination of both. As we found a pool of AP-1 colocalizing with E-cad at AJs, we focused on testing the potential role for AP-1 in E-cad internalization from the plasma membrane. We used Fluorescence Recovery After Photobleaching (FRAP) to measure the dynamics of the *ubi*::E-cad-GFP at the AJs (Fig 7A-B). E-cad recovery in FRAP is diffusion-uncoupled and best-described by a bi-exponential model with the slow component mediated by endocytic recycling [58–60]. We found that AP-1 depletion halved the halftime of the slow component’s recovery (Fig 7B). E-cad recovery following AP-1 depletion was best-fit by a single exponential model, likely as the slow halftime reduction made it impossible to confidently separate the components [61]. The halftime of a diffusion-uncoupled recovery depends only on the reaction off-rate (internalization rate in this case) [59,62]: smaller halftime means larger off-rate. Therefore, we conclude that E-cad internalization from AJs is elevated upon AP-1 knockdown, which is consistent with AP-1 acting on endocytosis at AJs.

**Figure 7.**
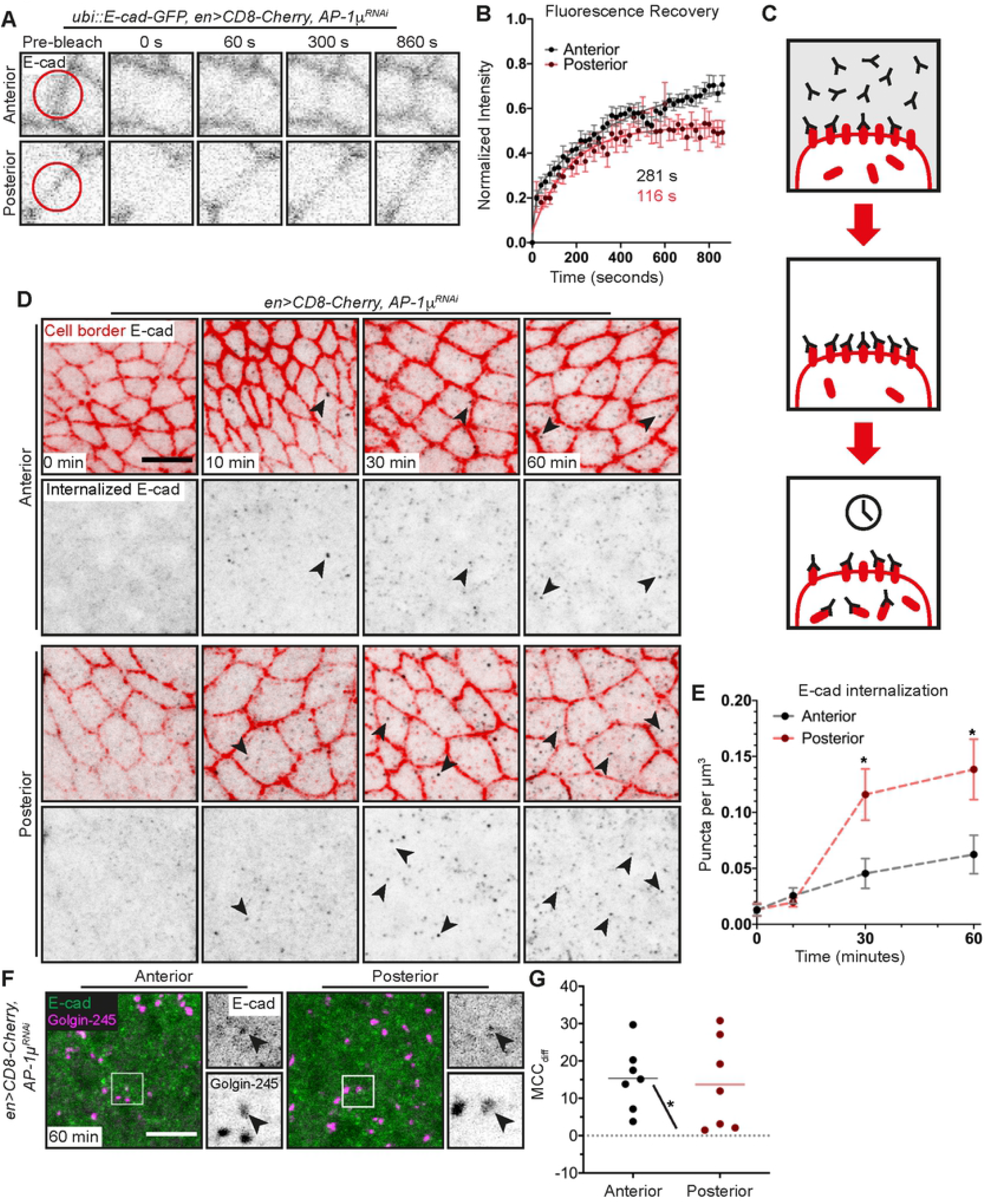
AP-1 decelerates E-cadherin endocytosis. (A) Frames of *ubi*::E-cad-GFP FRAP at wing disc cell borders at the indicated compartments and timepoints. Red circles in prebleached frame (P) outline the bleached spots. (B) Average recovery curves (mean ± s.e.m., n=16 and 15 cells, from 7 wing discs), with the best fit curves (solid lines) and slow halftimes. (C) Cartoon depicting the pulse-chase protocol to measure uptake of E-cadherin antibody. (D) Pulse-chase labelling with E-cad antibody in the anterior (control) and posterior (expressing AP-1μ RNAi) compartments of wing discs. Apical region with AJs is in red; the puncta (examples indicated with arrowheads) in the 1.9 μm below – in black. Scale bar: 5 μm. (E) The number of vesicles per μm^3^ during pulse-chase labelling (mean ± s.e.m., n=5-7 compartments/ time point). *P<0.05 (Welch’s t-test). (F) Pulse-chase labelling (60 minutes) with E-cad antibody (green, left) and stained for Golgin-245 (magenta, left), in the anterior (control) and posterior (expressing AP-1μ RNAi) compartments of a wing disc. Grayscale images correspond to E-cad (top) and Golgin-245 (bottom) signal within the white square on the left image. Scale bar: 5 μm. (G) Manders’ Correlation Coefficient difference (MCC_diff_, see Methods) for the co-localization between internalized E-cad antibody and Golgin-245. Each dot represents the average for individual paired compartments (n=7 discs). *P<0.05 (Wilcoxon signed-rank test).

Next, we validated the increased endocytic internalization of E-cad from the plasma membrane with the pulse-chase assay (Fig 7C). We measured the internalization of an antibody against the E-cad extracellular domain bound to E-cad at the cell surface as a readout of the E-cad endocytosis from the membrane (Fig 7D) [58]. In this instance, we measured E-cad-antibody-containing puncta just below the AJs belt to confidently exclude the signal coming from the plasma membrane while remaining within the area rich in the TGN and RE signal (Fig S4). We found that the number of vesicles containing E-cad antibody doubled following AP-1μ knockdown at 30 and 60 minutes after labelling (Fig 7E). As expected, this internalised E-cad co-localized with TGN both in the control and AP-1 depleted cells (Fig 7F-G).

We finally asked if the elevated cytoplasmic E-cad and its cytoplasm/AJs ratios were responsible for apoptosis following AP-1 knockdown (Fig 1H), as in the wing discs the failure to localize E-cad at AJs induces apoptosis through activation of JNK signalling [63]. We examined if inhibiting E-cad endocytosis ameliorated the phenotype, which we observed upon AP-1 knockdown. To specifically block endocytosis of E-cad without effects on general endocytic machinery, we overexpressed p120-catenin, which directly binds the E-cad intracellular domain and prevents its internalization in both flies and mammalian cells [59,64,65]. While overexpression of p120-catenin did not affect wing size by itself, co-expression of p120-catenin with AP-1μ RNAi increased the wing size relative to AP-1 knockdown indicating a partial rescue (Fig 8A, B). To determine the cause of this rescue, we examined cell death and integrin localization in wing discs co-expressing p120-catenin with AP-1μ RNAi. There was no difference in αPS1 integrin localization with or without p120-catenin overexpression in wing discs expressing AP-1μ RNAi (Fig 8C), which was consistent with wing blisters observed in wings coexpressing p120-catenin with AP-1μ RNAi (Fig 8A). However, there was less cell death visualised using Dcp-1 antibody in wing discs that overexpressed p120-catenin than in those that did not upon AP-1μ knockdown (Fig 8D, E). Therefore, we conclude that reducing E-cad endocytosis in AP-1 depleted cells prevents cell death, suggesting that elevated E-cad endocytosis contributed to cell death induction upon AP-1 downregulation (Fig 8F).

**Figure 8.**
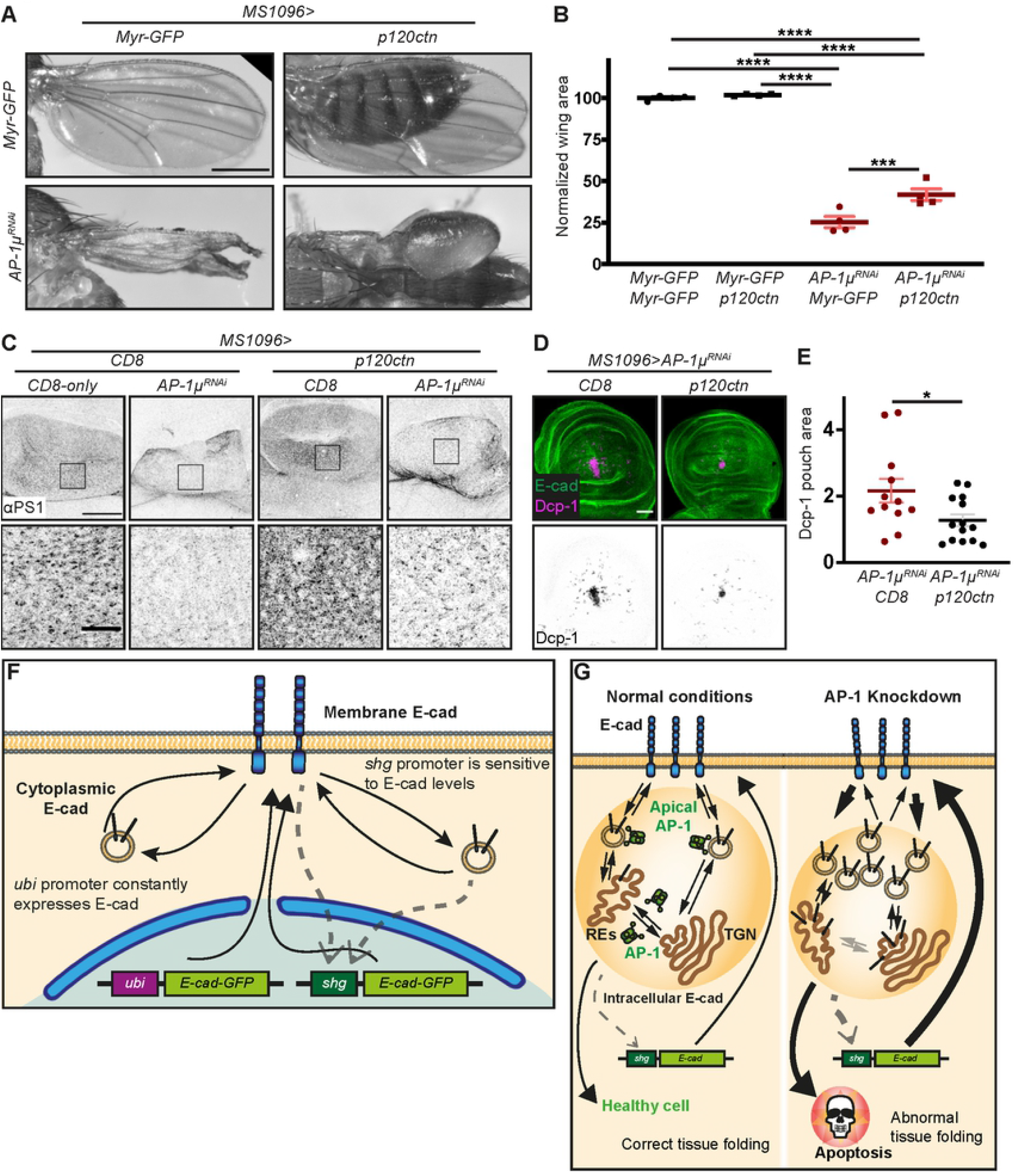
AP-1 promotes cell survival by controlling intracellular levels of E-cadherin and summary models. (A) Adult wings expressing the indicated proteins or AP-1μ RNAi in their wing pouches. Scale bar: 0.5 mm. (B) Relative area (normalized to Myr-GFP controls) of adult female wings depicted in A. Dots represent individual adults (n=4 per genotype). ***P<0.001 and ****P<0.0001 (Brown-Forsythe and Welch ANOVA test). (C) Basal region of wing discs expressing either CD8-Cherry or AP-1μ RNAi together with CD8-Cherry (a single copy in the case of CD8-Cherry) or p120ctn, stained for αPS1 (grayscale) with regions highlighted by rectangles shown in the bottom row. Scale bars: 50 μm (top); 10 μm (bottom). (D) Wing pouch regions of discs expressing AP-1μ RNAi with CD8-Cherry (left, not shown) or p120ctn (right) in the posterior compartment visualized with E-cad antibody (green, top) and cleaved caspase-3 (Dcp-1, magenta, top; inverted grayscale, bottom). Scale bar: 50 μm. (E) Percentage of wing pouch region occupied by Dcp-1 signal. Dots represent individual discs for each genotype (n=12 and 14). *P<0.05 (unpaired t-test). (F) Model of the proposed E-cad feedback. The cytoplasmic/AJs ratio of E-cad influences the transcription of the *shg* gene. Only the endogenous *shg*::E-cad-GFP is affected by this. (G) Model of the functions of AP-1 in the developing wing discs. In control (left), besides vesicle trafficking between the Recycling Endosomes (REs) and the Trans Golgi Network (TGN), a subapical fraction of AP-1 localizes at/near the AJs. Upon AP-1 knockdown (right), E-cad is retained in TGN/REs following its elevated endocytosis from the AJs and possibly delayed delivery to the AJs. This increase of E-cad intracellular levels induces cell death. To bring E-cad membrane levels back to normal, cells increase *shg* expression.

## Discussion

In this study, we report complex morphological changes caused by the reduced function of the AP-1 complex. While such a pleiotropic effect is expected from proteins with a general role in intracellular trafficking, the reported phenotypes are very specific and can be attributed to a limited number of specific cargoes, e.g. the ectopic folding is consistent with the defects in basal targeting of integrins. Concurrently, the elevated cell death can be attributed to hyperinternalization of E-cad from the cell surface as it was reduced by inhibiting E-cad endocytosis (Fig 8D-E). Such an effect of AP-1 downregulation on E-cad at the cell surface was rather unexpected, but do agree with the co-localization of a distinct subapical AP-1 pool with AJs that we report. Finally, we found that this hyperinternalization of E-cad was accompanied by increased E-cad expression, compensating for the loss of E-cad at the plasma membrane (Fig 4F and 8F).

While localization of AP-1 at AJs was not previously reported, it is not surprising as AP-1 has been found at the plasma membrane in such diverse systems as the cell-ECM adhesion in mammalian kidney cells and phagosomes in *Dictyostelium* amoebae [66,67]. Additionally, both Arf1 and Arf6 – two small GTPases, which contributed to the recruitment of AP-1A and AP-1B, respectively, in mammalian cells – are found at the plasma membrane, where they regulate endocytosis in both fly and mammalian cells [31,65,68–72]. Moreover, Arf1 and AP-1 share the localization pattern being present at both TGN and AJs. Their involvement in both endocytosis and delivery to the plasma membrane provides a potential mechanism for integration of these processes, whereby changes in one lead to fine-tuning of the other. The coupling of exocytosis and compensatory endocytosis is established in neuronal cells, where SNARE-proteins and Ca^2+^ concentrations adjust the balance between two processes. To understand the consequences of such coupling, it would be important to determine proteins, whose trafficking is affected by each pool of AP-1, and if both pools act on the same proteins. However, dissecting the functions of individual pools is challenging. For example, while we established that knockdown of AP-1 promotes E-cad endocytosis, we cannot exclude a concurrent decrease in E-cad removal from REs/TGN, as AP-1B is required for E-cad delivery to the plasma membrane in mammalian cells [73].

Intriguingly, E-cad endocytosis was enhanced upon the loss of AP-1, as it is a trafficking adaptor usually thought to promote vesicular transport. Similarly, in mammalian cells (LLC-PK1), expression of AP-1B slowed down cell migration which is consistent with the reduced integrin turnover [66,74]. One of the scenarios proposed for the AP-1B function at the plasma membrane in these cells was that AP-1B forms coats that abort without scission but at the same time occupy binding sites for AP-2 and thus reduce the net rate of AP-2-mediated endocytosis. Alternatively, both AP-1 and AP-2 may be involved in E-cad endocytosis, but clathrin-coated pits mature faster in the case of AP-2 recruitment due to the presence of specific modifying enzymes such as kinases, which can alter the speed of maturation [75]. Then in the absence of AP-1, the remaining AP-2 driven endocytosis would lead to the faster net rate of E-cad internalization. In either scenario, it makes physiological sense that AP-1 slows E-cad internalisation from AJs. If there is a limited common pool of AP-1, its reduced localization to TGN would potentially lead to increased AP-1 availability for recruitment to AJs. Such reduced localization of AP-1 to TGN could be a consequence of impairment in TGN function and a sign of reduced E-cad delivery to the plasma membrane. In this case, the concurrent elevation of AP-1 at AJs will reduce E-cad removal from the plasma membrane, thus restoring E-cad levels at AJs and contributing to adhesion robustness.

Such robustness of adhesion is further supported by our observation of increased *shg* expression upon AP-1 knockdown (Fig 4F), suggesting existence of a feedback loop. We have recently reported feedback between E-cad protein levels and its gene expression; specifically, the recruitment of the transcription regulator STAT92E to E-cad adhesion via the Par-3 protein limits its availability for binding the Heterochromatin Protein 1 (HP-1) in the nucleus [55]. This mechanism allows restoring E-cad expression when its levels are altered; elevated E-cad leads to reduced nuclear STAT92E and hence, HP-1 function, which not only reduces heterochromatin formation but also E-cad expression. Increased *shg* expression in cells lacking AP-1, suggests that this feedback mechanism might be activated not only by the total E-cad protein level but also by the enhanced internalization of E-cad from the cell surface and retention of this internalized E-cad at TGN/REs (Fig 8F).

In addition to promoting *shg* expression, we found that the relative increase of intracellular E-cad compared to the membrane pool induced cell death. Disrupted localization of E-cad to AJs in cell mutant for actin capping protein led to induction of JNK-mediated apoptosis, whereas inhibition of this apoptosis resulted in overproliferation [63]. It is worth mentioning that the loss of the capping protein also leads to increased E-cad expression, supporting the feedback mechanisms described above. Similarly, we observed hyperplastic overgrowth of wing discs when apoptosis was inhibited upon AP-1 knockdown (Fig 1H). Therefore, apoptosis induction following E-cad hyperinternalization may provide a protective mechanism, whereby defective cells are eliminated (Fig 8G). This mechanism agrees well with the multistage theory of cancerogenesis [76], so that additional sequential mutations are required to overcome both this cell-death inducing mechanism and the feedback which restores E-cad levels.

In summary, we provide evidence that a subapical pool of the AP-1 complex exerts a brake upon E-cad internalization, excess of which promotes cell death. We demonstrate that this AP-1 function is accompanied by transcriptional feedback which maintains E-cad levels at the cell surface. In line with the correlation of the AP-1B loss with the metastatic potential in humans [66], our findings highlight the pivotal role of the AP-1 complex in preventing tumour progression and enabling correct epithelial morphogenesis.

## Materials and Methods

### Reagents and tool table

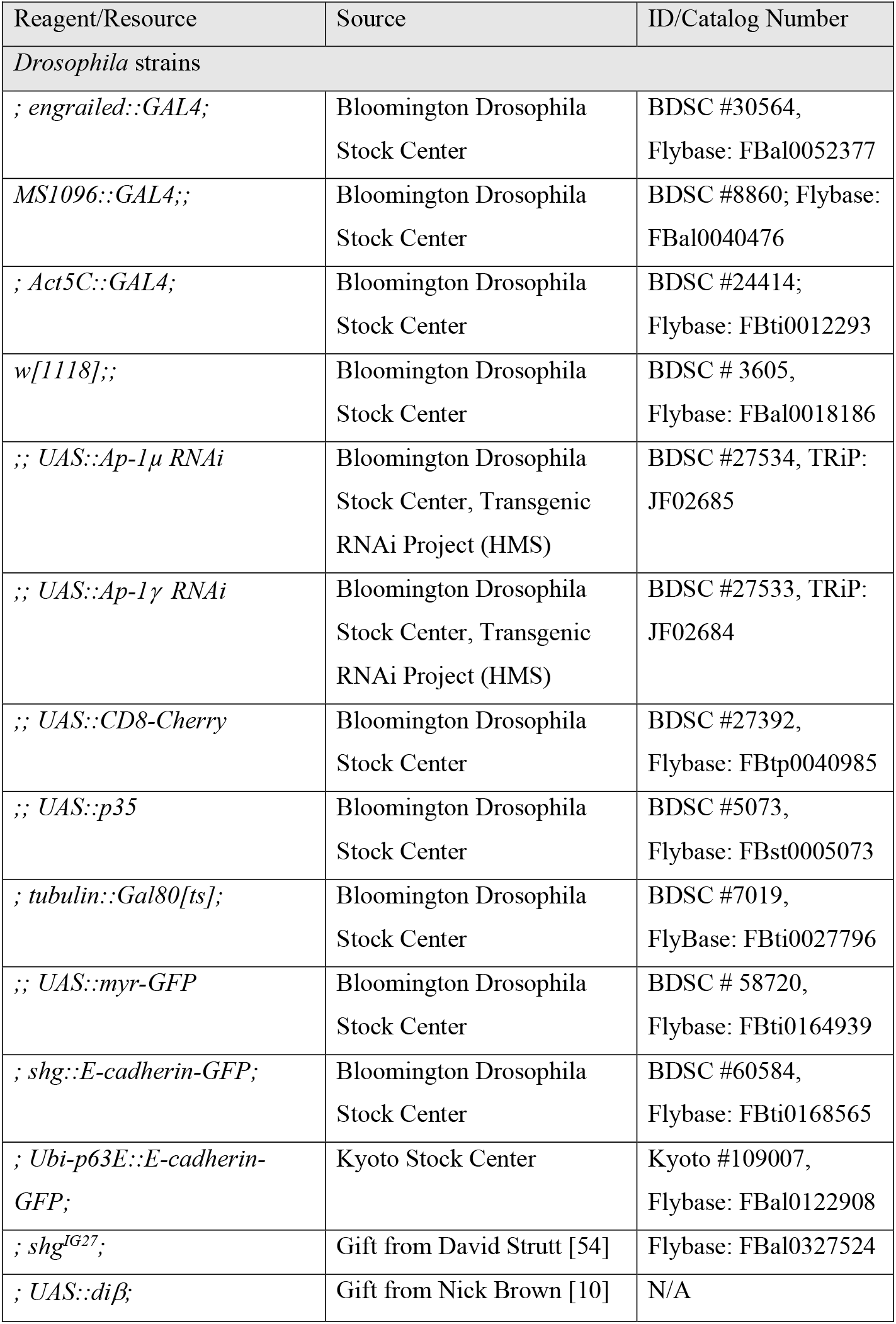
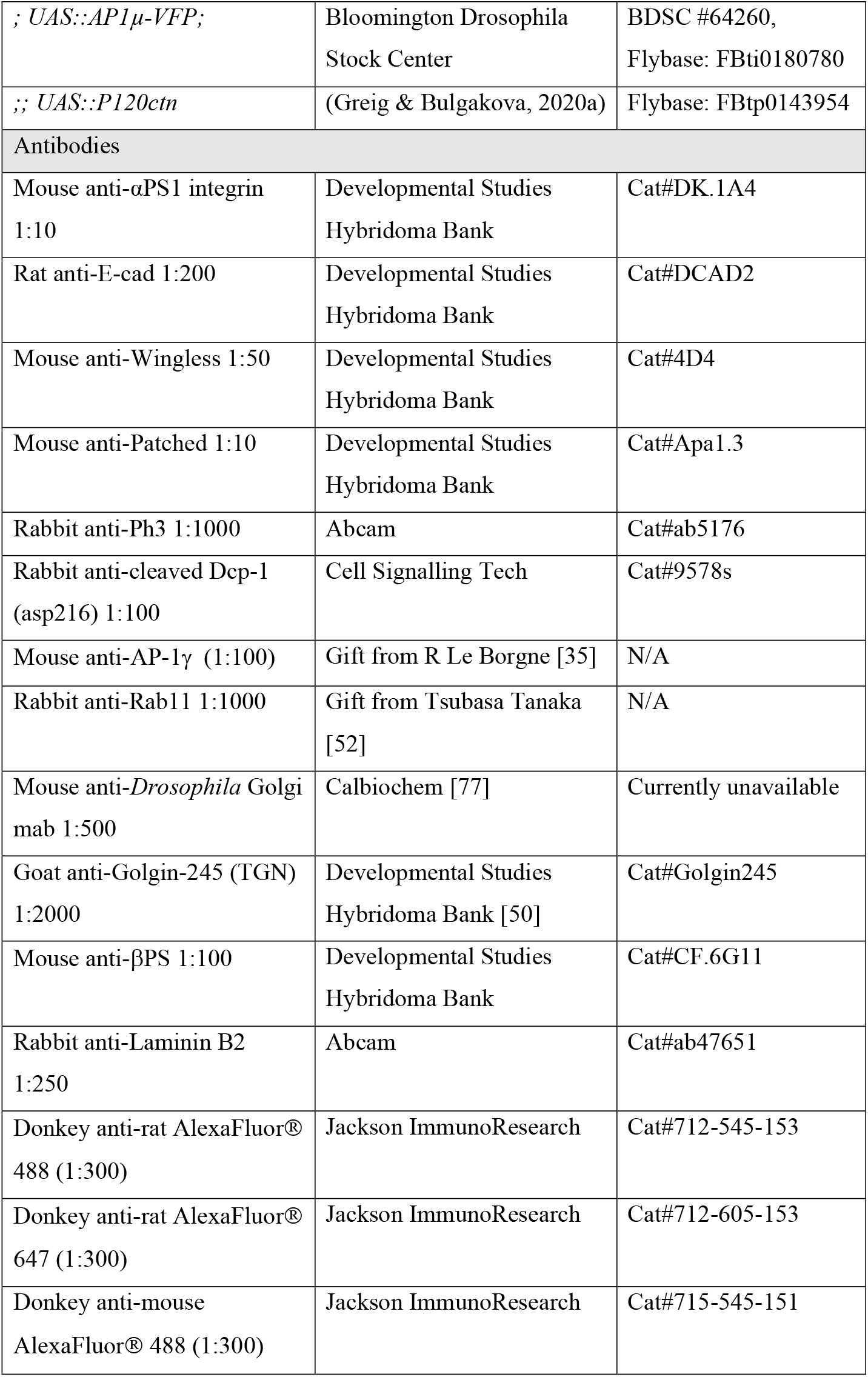
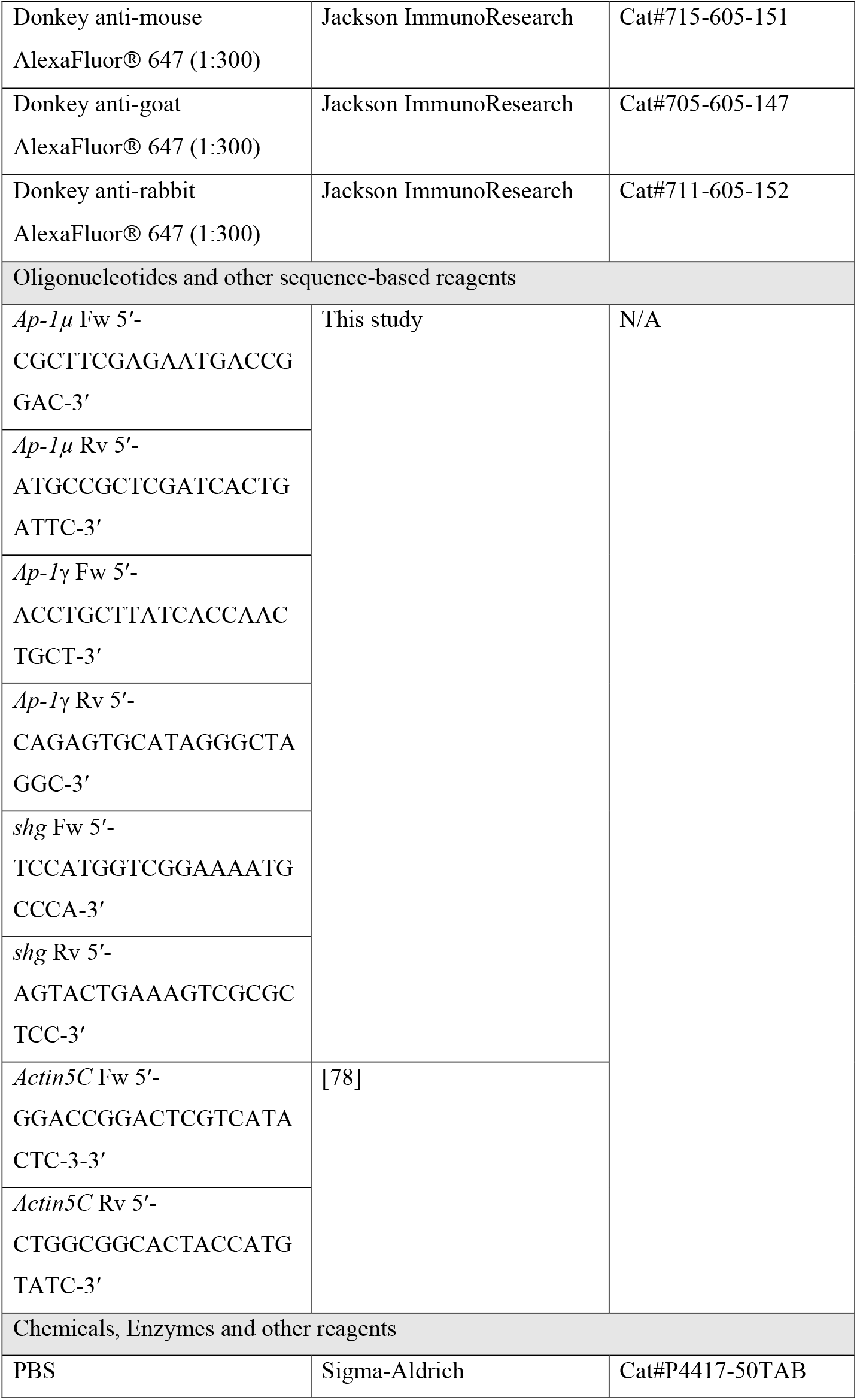
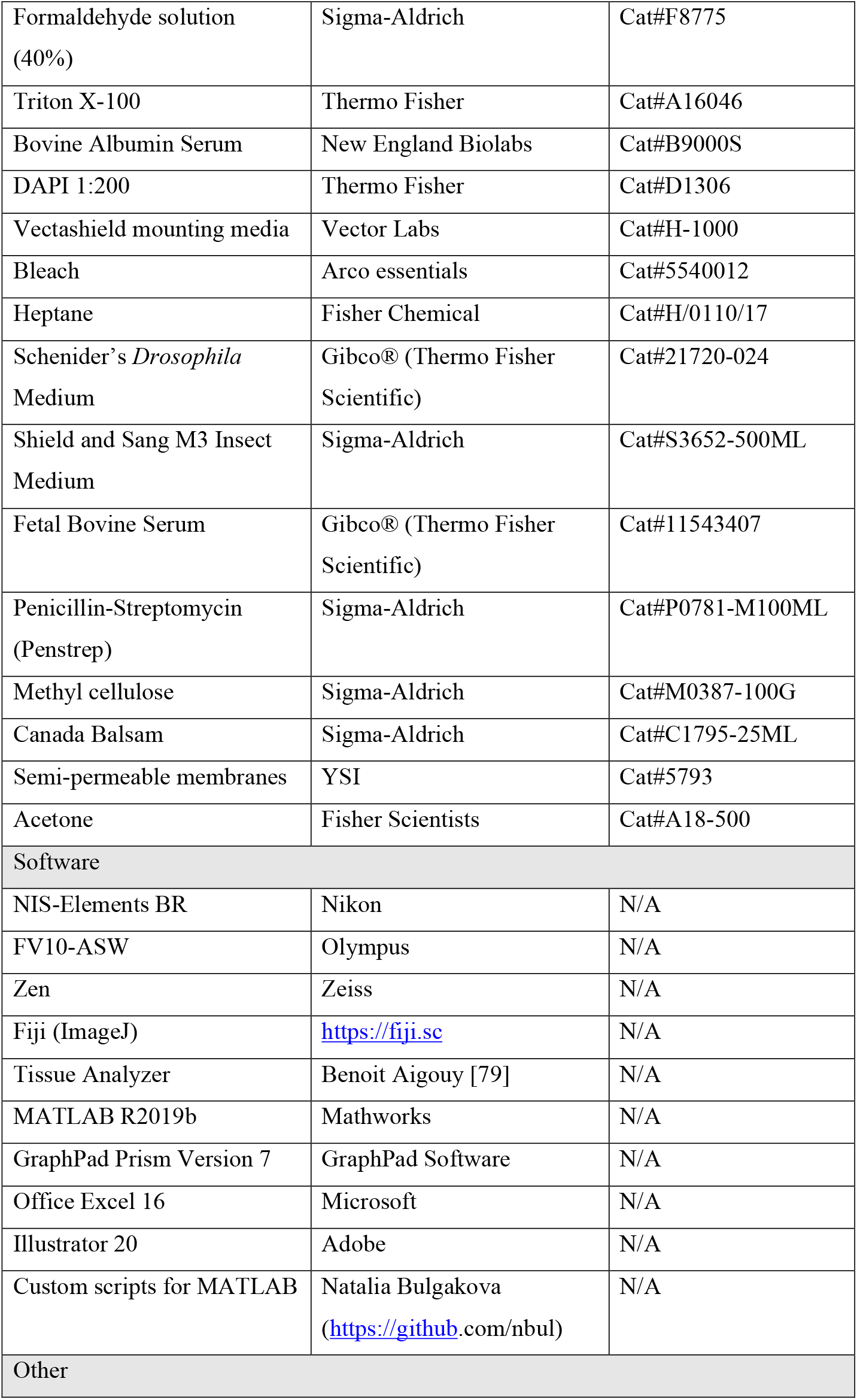
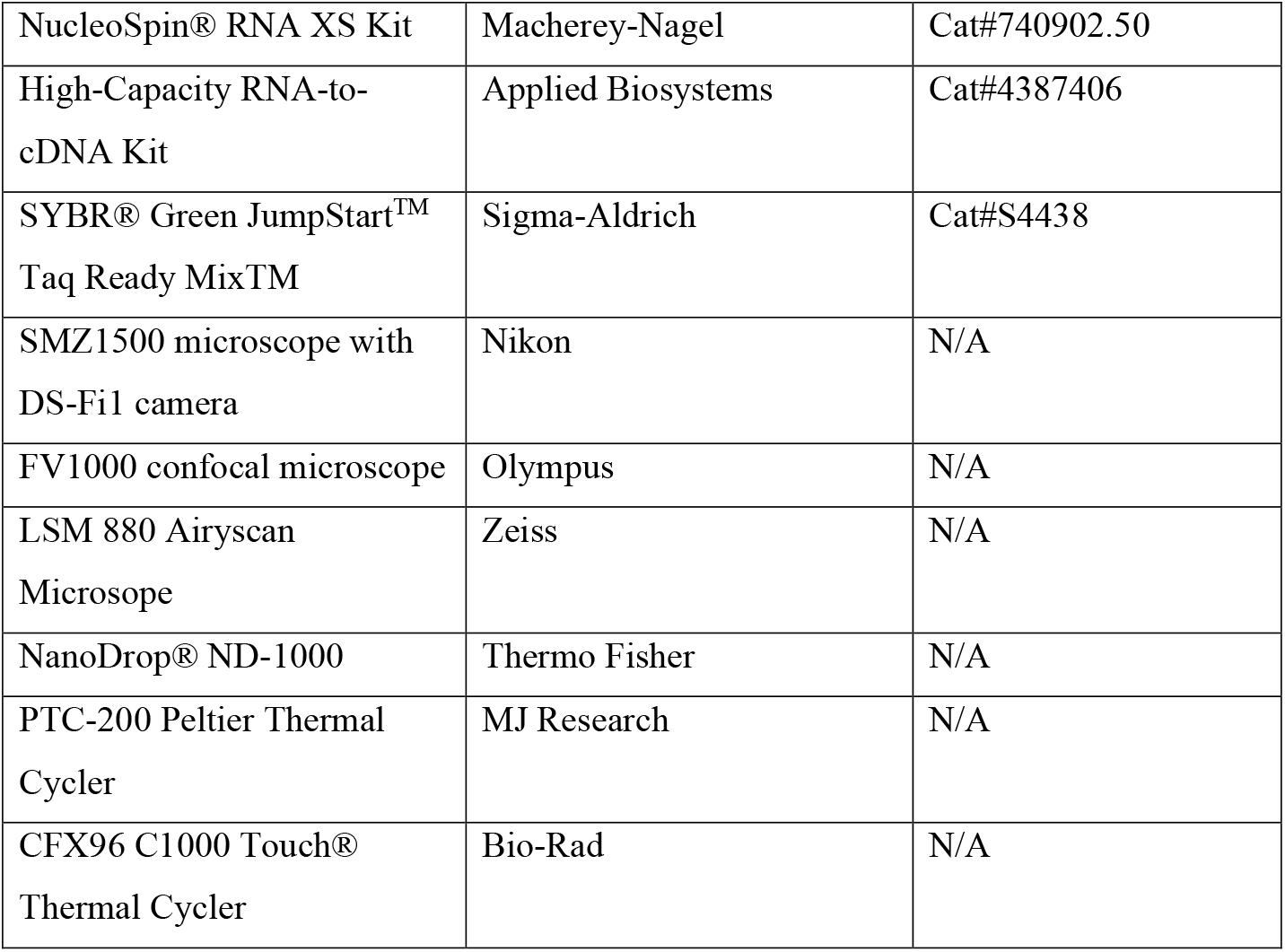

### Fly genetics and husbandry

*Drosophila melanogaster* were raised on standard cornmeal/agar/molasses media at 18°C or 25°C unless otherwise specified. To express constructs of interest, the GAL4/UAS system was used [40], with *engrailed*::GAL4 (*en*::Gal4), *MS1096*::GAL4, or *Act5C*::GAL4 drivers. To examine adult wings, flies were raised at 18°C to circumvent potential lethality due to RNAi expression. To study larval wing discs, larvae were raised at 25°C. The GAL4:UAS ratio was kept constant within each experimental dataset using additional copies of *UAS*::CD8-Cherry or *UAS*::myristoylated-GFP (*UAS*::myr-GFP). Acute expression of RNAi was achieved with the combination of *Act5C*::GAL4 and *tubulin*::GAL80^ts^. Larvae then were raised at 18°C for thirteen days after egg laying and shifted to 29°C for 48 hours prior to dissection of the wing discs.

### Adult wings imaging

Female adult flies were frozen upon collection with CO_2_, with the wings removed for direct imaging. For the quantification, frozen female flies were dehydrated in serial solutions of EtOH (50%, 70% x2, 90% x2) for 10 minutes each, followed by 10 minutes in acetone. Flies were kept overnight in fresh acetone and the wings were then mounted in Canada Balsam (Sigma, C1795-25ML). Wings, or representative adult females, were imaged with a Nikon SMZ1500 microscope equipped with Nikon DS-Fi1 camera controlled by NIS-Elements BR software.

### Dissection and Immunostaining

#### Wing discs

Third instar larvae were dissected after being kept for six days after egg laying at 25°C. Cuticles with attached imaginal discs were fixed for 15 minutes with 4% formaldehyde (Sigma, F8775) in PBS (Phosphate Buffer Saline, Sigma-Aldrich, P4417) at room temperature, then washed with PBS with 0.1% Triton X-100 (Thermo Fisher, A16046, hereafter PBST), and incubated with 1% Bovine Serum Albumin (BSA, New England Biolabs, B9000S) in PBST for 1 hour at room temperature. Cuticles were incubated with primary antibodies and 1% BSA in PBST overnight at 4°C, washed in PBST, incubated overnight at 4°C with secondary antibodies and 1% BSA in PBST (including DAPI when applicable), and washed in PBST again. All antibodies and their concentrations are listed in the Reagents and Tools Table. Finally, discs were separated from the cuticles and mounted in Vectashield (Vector Labs, H-1000). For samples not requiring immunostaining, e.g. imaging direct fluorescence of E-cad-GFP, discs were immediately mounted following fixation and one round of washing in PBST.

#### Embryos and retinas (Fig S2)

*Drosophila* embryos were aged up to the desired developmental stage with 3-hour collections at 25°C, followed by 21 hours at 18°C. Embryos were then dechorionated in 50% commercial bleach solution in water for four minutes. Following extensive washing with deionised water, embryos were fixed with a 1:1 4% formaldehyde in PBS : heptane (Fisher Chemical, H-0110-17) for 20 minutes with constant agitation at room temperature. Embryos were devitellinized by vigorous shaking in 1:1 methanol : heptane for 20 seconds, washed and stored in methanol at -20°C upon required. Methanol was removed by washing in PBST, and embryos were subjected to the same staining procedure as the wing discs but using 0.05% Triton in PBS rather than 0.1%.

Prepupal stage individuals raised at 25°C were collected and aged for 40 hours at 25°C to reach the desired developmental stage. Retinas were extracted by cutting the external cuticle open in PBS. The same protocol as for wing discs was followed for staining.

### Pulse-chase assay

Wing discs were dissected in Schneider’s Insect medium (Gibco, 21720-024) with 1% Penstrep (Sigma-Aldrich, P0781-M100ML) and 5% Fetal Bovine Serum (FBS, Gibco, 11543407), and the peripodial membrane was removed with a fine needle. Then, the discs were incubated with rat anti-E-cad antibody (DCAD2, 1:200, DSHB) in the same medium at 4°C for 1 hour. After three quick washes with antibody-free cold medium, the discs were incubated in fresh antibody-free medium at room temperature for 10, 30 or 60 minutes. Several (5-7) discs were fixed in 4% formaldehyde in PBS for 15 minutes at each analysed time point (immediately after washing for t = 0 min). The discs were stained as described above using AlexaFluor® 488-conjugated anti-rat antibody (Jackson ImmunoResearch).

### FRAP

Wing discs were dissected in Shield and Sang M3 Insect Medium (Sigma, S3652-500ml) with 1% Penstrep (Sigma-Aldrich, P0781-M100ML) and 5% Fetal Bovine Serum (FBS, Gibco, 11543407), and placed into an observation chamber with a poly-D-lysine coated coverslip [80]. Then, the medium was replaced by Shield and Sang M3 Insect Medium supplemented with methyl-cellulose (Sigma, M0387-100G). Discs were positioned carefully to adhere to the coverslip and covered with an oxygen semipermeable membrane (YSY 5793).

### Microscopy

All microscopy experiments except colocalization analyses were done using an upright Olympus FV1000 confocal microscope with either 60x/1.40 NA (fluorescence intensity, FRAP, and cell morphology) or 20x/0.75 NA (tissue morphology) objectives. In the former case, images were taken in the dorsal region of the wing disc pouch (Fig 1B): for each disc and compartment a z-stack of 6 slices spaced by 0.38 μm was acquired capturing the complete span of the apical Adherens Junctions. This z-spacing was used for all the acquisitions with the 60x objective, while 1 μm spacing was used for z-stacks done with the 20x objective. All the images were at 16-bit depth in Olympus binary image format. For FRAP, several circular regions of 1 μm radius were photobleached at junctions so that there was only one bleach event per cell. Photobleaching was performed with 8 scans at 4 μs/pixel at 80% 405 laser power, resulting in the reduction of E-cad–GFP signal by 60–80%. Z-stacks were acquired just before photobleaching, and immediately after photobleaching, and then at 20-s intervals for 15 min in total.

Images used for colocalization assays were taken using an inverted Zeiss LSM 880 Airyscan confocal microscope. Z-stacks of 20 slices spaced by 0.20 μm (AP-1μ-VFP co-localization) or 10 slices spaced by 0.18 μm (E-cad GFP co-localization) were taken with a 63x/1.40 NA objective. Raw 16-bit images were processed using Zeiss software (automatic mode) to obtain “.czi” files.

### Image processing

#### Compartment size (wing discs and adult wings)

Z-stacks with E-cad-GFP and CD8-Cherry signals were projected using the “maximum intensity” algorithm in Fiji (https://fiji.sc) and used to measure the area of the whole disc and posterior compartment, respectively, with the Fiji selection tool. The anterior compartment area was calculated by subtracting the posterior compartment area from that of the disc. For adult wings, posterior compartment was measured with the Fiji selection tool using the vein L4 as its border, while the vein L3 acted as the border of the anterior compartment, leaving out the area between both veins.

#### Apical cell area, membrane and cytoplasmic protein levels

For the analyses of membrane and cytoplasmic protein levels, z-stacks with signal of GFP fluorescence were projected using the “average intensity” algorithm in Fiji. To distinguish membrane and cytoplasm as well as measure apical cell area, we generated binary masks from images with visualized cell outlines (i.e. E-cad signal). In particular, z-stacks with E-cad signal were projected using the “maximum intensity” algorithm in Fiji. The background was subtracted from these projections using a rolling ball of 50-pixel radius, and their brightness and contrast were adjusted using the automatic optimization algorithm in Fiji. These projections were used to generate masks using the Tissue Analyzer plugin in Fiji [79]. Cells in which the AJs were not completely in focus were removed manually.

The binary masks were used to measure the fluorescence intensity of average projections using our in-house Matlab script (https://github.com/nbul/Intensity). First, individual objects (cells) and their boundaries were identified from each binary mask. Then the identified objects (cells) were analysed on a cell-by-cell basis. The area of objects and length of their boundaries in pixels were determined, and then manually converted from pixels to μm^2^. The boundary was dilated using a diamond-shaped morphological structural element of size 3 to encompass the XY spread of E-cad signal. The mean and total (sum of all pixel intensities) intensities of the dilated boundary (membrane signal) and the object with subtracted boundary (cytoplasm) were calculated. All the values were averaged to produce single values per wing disc thus testing biological replicates and excluding chances of one disc having higher contribution to the result due to variable number of cells in each disc.

#### Proliferation

z-stacks with E-cad-GFP and pH3 (647) signal corresponding to the anterior and posterior dorsal wing pouches were processed separately for each channel. E-cad signal was segmented as described above using Tissue Analyzer. The resulting binary masks were dilated and then inverted. The area with cells was used then to generate maximum projections of the equivalent region for the pH3 channel. The projection was processed using the following steps: background was subtracted using a rolling ball of 1-pixel radius; then Gaussian blur of 2-pixel radius was added; and image was binarized using a threshold set to three times the average intensity of the original maximum projection.

To calculate the cell number and the total cell area in the imaged regions from the processed binary segmented masks, we employed the Particle Analysis plugin on Fiji, limiting particle-detection to sizes up to 67.5 um^2^ (or 3000 pixel^2^). The number of proliferating cells in the same area was determined using the processed images of the pH3 channel corresponding to the same area using particle sizes between 5 and 50 um^2^. These numbers were used to calculate the number of dividing cells per 100 cells or per 100 um^2^.

#### Co-localization

For co-localization of AP-1μ-VFP with cellular markers as well as E-cad-GFP with AP-1γ antibody, we employed the co-localization plugin Coloc 2 in Fiji (https://imagej.net/Coloc2). We selected Manders’s Colocalization Coefficients (MCC [81,82]) as more informative for probes distributed to more than one compartment than Pearson’s correlation coefficients [83]. The MCC_diff_ value was obtained by subtracting the percentage of pixels positive for AP-1μ-VFP (obtained using the threshold determined by Coloc 2) from the percentage of pixels positive for the marker that were also positive for AP-1μ-VFP.

For co-localization of E-cad-GFP with Golgi bodies and recycling endosomes, we analysed the images with an in-house script at MATLAB (https://github.com/nbul/Localization). This script followed the same principle to obtain MCC_diff_ values as the Coloc 2 plugin, but is more versatile as it enables a manual selection of a threshold method for each probe. Such selection is required as the bright signal of E-cad-GFP at cell borders limits its detection in cytoplasm using standard threshold methods. The E-cad-GFP signal at cell borders was excluded from the analysis, and the resulting MCC_diff_ values accounted only therefore for the intracellular signal.

#### Pulse-chase

12 sections, comprising 6 sections spanning the complete region of Adherens Junctions visualized with the staining of the antibody bound to E-cad at the cell surface and the 6 sections immediately basal to Adherens Junctions, were used for counting the E-cad-positive vesicles. Firstly, a binary mask with cell outlines was created from a projection of the 6 apical-most sections as described above. The average cytoplasmic signal and cell area were measured with our in-house script as described above and used to threshold the corresponding maximum projection of the 6 basal sections in Fiji using the following steps: background was subtracted using a rolling ball of 4-pixel radius; then Gaussian blur of 2-pixel radius was added; and image was binarized using a threshold set to 70% of the average cytoplasmic signal of the apical section. The resulting images were used for the analysis using the Particle Analysis plugin in Fiji, limiting particle-detection to sizes between 0.01 and 0.45 μm^2^. The parameters were empirically selected through interactive validation. Puncta density was determined using the average apical area of the cells.

#### FRAP

z-stacks for each time point in the time-series were projected using the “average intensity” algorithm in Fiji. The recovery curves were obtained by manually measuring intensities of background, control region, and photobleached region using 2-μm diameter circular regions for each time point in Fiji. The raw data was processed as in [61] to obtain the normalized recovery curves, namely following background subtraction the intensity at the bleached spot was normalized to the intensities of a control area at the same time point and the bleached area before bleaching.

#### Cell death

z-stacks of six slices corresponding to the basal region of the discs were averagely projected in Fiji. E-cad antibody signal was used to generate a mask encompassing the whole disc pouch (all the disc ventral to the H/H fold, corresponding approximately to the domain of *MS1096* expression. Dcp-1 signal was processed twice with the “smoothening” tool and then thresholded with Otsu method (automatic levels). Dcp-1 signal was measured as a percentage within the mask.

#### Other processing

Sagittal views of the wing discs, for example in Fig 1, were generated using the “Reslice” tool in Fiji. For representative cases shown in Figures, maximum projections of the regions of interest were generated in Fiji using minimum modification, such as tilting, cropping, or automatic contrast of the whole view, for better presentation.

### RQ-qPCR

Primers for RT-qPCR were designed and *in silico* tested using Flybase (https://flybase.org), Primer-BLAST (https://www.ncbi.nlm.nih.gov/tools/primer-blast/index.cgi), and Net Primer (http://www.premierbiosoft.com/netprimer/) tools, and whenever possible aimed to target sequences separated by introns and present in all the splicing variants. All primers and amplicon sizes are listed in the Resources and Tools Table, and were manufactured by Thermo Fisher. RNA was extracted from wing discs dissected on ice using the NucleoSpin® RNA XS Kit (Macherey-Nagel, 740902.50). Control wing discs expressed *UAS*::myr-GFP instead of the RNAi. For AP-1γ knockdown, only discs from male larvae were used to exclude the potential effects of dosage compensation. RNA concentration was determined with an ND-1000 Spectrophotometer (Nanodrop®, Thermo Fisher) and immediately used to generate cDNA with the High-Capacity RNA-to-cDNA Kit (Applied Biosystems, 4387406) using 400 ng of RNA. cDNA was then used as a template for the quantitative PCR reaction using SYBR® Green JumpStart™ Taq Ready Mix™ (Sigma-Aldrich, S4438) using a CFX96™ Real-Time System (Bio-rad) and its proprietary software. All the primers were tested with standard reaction curves and gel electrophoresis. Upon testing of several widely employed housekeeping genes in *Drosophila* [84,85], primers against *Actin5C* were selected due to their reproducible performance. Reactions were carried out in 3 technical replicates per a biological replicate (15 wing discs), in a volume of 10 μl with a primer concentration of 1 pmol/μl. At least three biological replicates were done per genotype. Ct values were obtained from SYBR fluorescence using thresholds determined from the standard curves [86]. Primer purity was tested on a control without any template in every performed assay.

Expression levels were determined by the 2^-ΔΔCT^ method [87]. For each biological replicate, Ct values were averaged across all technical replicates; and the average Ct value of *Actin5C* was subtracted from the target gene. This result (ΔCT) was normalized by subtracting the average ΔCT of the control genotype, producing ΔΔCT. This value was converted using the formula 2^-ΔΔCT^.

### Statistical Analysis

All the statistical analyses were done in GraphPad Prism 7 (https://graphpad.com/scientific-software/prism/). First, the datasets were confirmed to be free of detectable outliers using the ROUT detection method (Q=0.1%), and the distributions were tested for being normal with D’Agostino & Pearson test. Precise n numbers, the type of statistical test, and the type of represented data (i.e. individual cases, Mean and SD, etc…) are described in the Figure legends. Significance was visually depicted in all the graphs with either the precise value or asterisks (* - p<0.05, ** - p<0.001, *** - p<0.0001, and **** - p<0.00001). For commonly used tests, such as t-test, two-tailed versions were used. Non-parametric tests were used when at least one sample did not display normal distribution, and appropriate corrections were applied if the assumption of equality of standard deviations was not met.

#### Posterior:anterior compartment size ratio

Posterior:anterior compartment size ratios of discs expressing RNAi against subunits of the AP-1 complex were tested against the external control discs expressing CD8-Cherry only using the Brown-Forsythe and Welch ANOVA test.

#### Proliferating cells

The number of dividing cells per 100 cells or per 100 um^2^ were compared with paired t-test/Wilcoxon test (between compartments of the same genotype) and Kruskal-Wallis test (between genotypes and the anterior control compartment).

#### Protein levels and cell size

Differences between genotypes were tested with One-Way ANOVA (normal distribution) or Kluskal-Wallis test. Differences between the paired sets of anterior and posterior compartments were tested with paired t-test (normal distribution) or Wilcoxon test.

#### Co-localization

The differences between control and experimental compartments were tested using paired t-test (normal distribution) or Wilcoxon test. The MCC_diff_ value was also tested against zero (random colocalization [82]) using the one sample t-test.

#### Pulse-chase

The puncta density was compared between the compartments using Welch’s t-test.

#### FRAP

Each replicate value in each dataset was considered as an individual point for curve fitting. GraphPad Prism software was used for nonlinear fitting using a bi-exponential model: f(t) = 1 − F_im_ − A_1_ × e^−t/Tfast^ − A_2_ × e^−t/Tslow^, and a single exponential model: f(t) = 1 − F_im_ − A_1_ × e^−t/Tslow^, where F_im_ is a size of the immobile pool of the protein, T_fast_ and T_slow_ are the half times, and A_1_ and A_2_ are the amplitudes of the fast and slow components of the recovery. An F-test was used to choose the equation and compare datasets.

#### Expression analyses

The gene expression in RNAi-expressing samples was compared to myr-GFP-expressing controls using Welch’s t-test.

#### Dcp-1 area

Dcp-1-positive wing pouch surface was compared between genotypes using unpaired t-test.

## Data availability

The generated *Drosophila* strains are available upon request. All in-house scripts are available at https://github.com/nbul/

## Acknowledgements

We thank Nick Brown, Iwan Evans, Roland Le Borgne and Tsubasa Tanaka for reagents; the University of Sheffield’s Wolfson Light Microscopy and Fly Facilities for their technical support; and Kyra Campbell and David Strutt for reagents and critical feedback on the manuscript.

